# Preconceptional immunomodulation partially corrects pregnancy abnormalities induced by endometriosis in a mouse model, with normalization of transcriptional alterations observed in the developing fetal-maternal interface at the single cell level

**DOI:** 10.64898/2026.01.02.695166

**Authors:** Kheira Bouzid, Roxane Bartkowski, Alix Silvert, Fabiana Moresi, Camille Souchet, Marine Thomas, Isabelle Lagoutte, Vaarany Karunanithy, Brigitte Izac, Charles Chapron, Pietro Santulli, Frédéric Batteux, Céline Mehats, Louis Marcellin, Ludivine Doridot

**Affiliations:** Université Paris Cité, CNRS, Inserm, Institut Cochin, F-75014 Paris, France; Department of Gynecology Obstetrics II and Reproductive Medicine, AP-HP, Hôpital Universitaire Paris Centre, Centre Hospitalier Universitaire Cochin, F-75014 Paris, France; Université Paris Cité, iPOP-UP

## Abstract

Endometriosis is a chronic disease of gynaecological origin that affects approximately 10% of women worldwide and has a significant impact on patients’ lives. Women with endometriosis attempting to conceive may experience infertility and have a higher risk of obstetric complications. They are also more likely to miscarry and have placental defects during pregnancy. The impact of endometriosis on pregnancy remains unclear. Defects in implantation and placentation may be present, but these processes are difficult to study in human subjects. To address this issue, we used a surgically induced mouse model of endometriosis and examined pregnancy in the CBAxDBA crossbreed, which is commonly employed to study pregnancy immunology. We examined early (E9.5) and late (E18.5) gestational stages.

In this model, we confirmed that the presence of endometriosis-like lesions resulted in fewer implantations and a higher proportion of fetal resorption, which is analogous to miscarriage. When an immune tolerance was induced previous to the endometriosis induction, we observed that endometriotic lesions were smaller, and gestational complications were mostly corrected. At early gestation (E9.5), fetal-maternal interfaces were retrieved to assess the potential impact of endometriosis and immunomodulation on placental development using single-cell RNA-sequencing. Our data show that most changes induced by endometriosis occur in decidual stromal cells, which originate from the endometrium. In these cells, we observed a consistent upregulation of Gata4, a transcription factor previously found to be elevated in the endometrium of women with endometriosis. We also observed downregulation of Prap1, which is an indicator of uterine receptivity and successful implantation in mice. In the mice where immune tolerance was induced, the endometriosis associated transcriptomic changes were partially reversed. In immune cells, endometriosis leads to an inflammation-associated signature. Additionally, the interferon gamma response is reduced in NK cells, which has been shown to be crucial for spiral artery remodelling and decidual integrity, which may be involved in the observed increased resorption rate.

Overall, our findings provide new insights into endometriosis-associated infertility and pregnancy complications in a mouse model that can be partially corrected by immune modulation. These results suggest that targeting the immune system should be considered in future studies to improve pregnancy outcomes in patients with endometriosis.

## Introduction

Endometriosis is a complex gynecological disease that affects approximately 10% of women of reproductive age. It is characterized by the presence of endometrium-like tissue in ectopic regions^1^. It is associated with pain, infertility, and a higher risk of pregnancy complications, significantly impacting patients’ lives.

Several studies have assessed the association between endometriosis and infertility and adverse pregnancy outcomes^2,3^. *In vitro* fertilization studies suggest that women with severe endometriosis have complications related to oocyte and embryo quality and difficult implantation^4^. In addition, women with endometriosis have an increased risk of miscarriage (odds ratio of 1.81 in spontaneous conception^5^) and other pregnancy complications, such as preeclampsia and *placenta previa*. At the end of pregnancy, a lower birth weight is found in babies born to endometriotic mothers^6–8^. These complications suggest that oogenesis, implantation, and placentation processes may be defective in endometriosis; however, the underlying mechanisms remain to be elucidated.

Several teams have developed mouse models of endometriosis, which are obtained either by grafting pieces of uterine horns on different tissues within the peritoneal cavity of the mouse by laparotomy^9–11^, or by cutting uterine horns/endometrium into small pieces and injecting them into the peritoneal cavity through an opening on the abdominal wall^12–14^. These studies have shown fertility and gestation complications in these models, with delayed coitus^14^, a lower pregnancy rate^9,10,12^, a reduced number of embryos^12^, a higher resorption rate^11–13^, and a lower pup weight^13^. These models can be used to study the underlying mechanisms of pregnancy outcomes in patients with endometriosis. In our team, we use a mouse model of endometriosis that we obtain by grafting uterine horn fragments on the peritoneum, which allows us to follow the lesion size using ultrasound. We have already shown in this model that peritoneal macrophages present inflammation induced by the presence of endometriosis lesions^15^. We can use this model to study the key players in the development of gestation, involving decidual and immune cells in the maternal part and trophoblast cells in the fetal part.

At the maternal-fetal interface early during gestation, most immune cells are innate immune cells (decidual Natural Killer cells – dNK and macrophages). They participate in fetal immune tolerance and implantation-placentation through the remodelling of the local tissue^16^. Women with endometriosis show altered immune cell phenotypes, such as low cytotoxicity in peritoneal NK cells^17^ and peritoneal macrophages with reduced phagocytic capacity and are prone to secrete pro-inflammatory cytokines^18^. Poor pregnancy outcomes in patients may be due to impaired immune cell function and recruitment at the maternal-fetal interface.

Interestingly, recent research has highlighted the concept of « trained immunity », where innate immune cells can develop a form of memory through exposure to bacterial by-products^3^, influencing their function in the context of inflammation^19^ and endometriosis^15^. Indeed, we recently showed that LPS^low^-trained immunomodulated macrophages could attenuate endometriosis in a mouse model obtained by grafting uterine fragments onto the peritoneal wall. Furthermore, coculture of human endometriotic cells with human LPS^low^-trained macrophages could reduce the fibro-inflammatory phenotype by inducing the downregulation of genes involved in cell adhesion and fibrosis^15^. LPS^low^ *in vivo* immune training can induce epigenetic and metabolic changes in innate immune cells, with a consequential change in gene expression, leading to an anti-inflammatory prone phenotype, mainly demonstrated in macrophages^19^, but similar innate memory mechanisms are described in NK cells^20^. This prompted us to investigate the potential benefits of LPSlow immunomodulation, specifically in the context of endometriosis-associated pregnancy complications. We used the abortion-prone CBA/J × DBA/2N mouse model, a well-established model for studying the general immunology of pregnancy^21^.

We first validated that we could recapitulate endometriosis-associated pregnancy complications in the CBA/J × DBA/2 mouse model. To determine how endometriosis affects the maternal-fetal interface, we generated and analyzed the transcriptome at the single-cell level of whole developing mouse placentas and isolated immune cells from these developing placentas with or without endometriosis. We demonstrated the beneficial effect of pre-conceptional LPS^low^ immunomodulation (IMM) on endometriosis-associated pregnancy complications in mice. By studying the transcriptome within the maternal-fetal interface, this study aimed to elucidate the mechanisms underlying endometriosis-associated pregnancy complications and the beneficial effects of pre-conceptional LPSlow immunomodulation.

## Material and methods

### Mice

Seven-to eight-week-old CBA/J female mice were purchased from Janvier Laboratory and housed under standard conditions (ad libitum food and water, 12h light-dark cycle) in the animal care facility of Institut Cochin.

### Endometriosis model

The endometriosis model consisted of a syngeneic graft of uterine horns to generate endometriosis-like lesions. In details, donor mice were sacrificed by cervical dislocation, and the uterine horns were surgically extracted and transferred to a Petri dish containing PBS. The uterine horns were opened longitudinally with micro scissors and 3 to 5-mm-length samples were prepared for grafting on the internal face of the peritoneum of the recipient mice (each horn fragment was weighed). A preoperative gavage of all donor mice with 100µg/kg/day of 17β-estradiol was performed for two days before sacrifice (to synchronize estral cycles). Recipient mice were anesthetized using isoflurane. An incision was made on the ventral midline, and one or two donor horn fragments were sutured onto the parietal peritoneum with two 7/0 polypropylene stitches (Prolen®, Ethicon, Somerville, NJ, USA). In all mice, tissue samples were sutured at identical positions in the abdominal wall to ensure that the host tissue sites exhibited comparable vascularization. The incision was then sutured with a 6/0 nylon thread.

### Immune training and endometriosis

Three independent experiments were performed using 5–10 mice per group using the previously described endometriosis model. The control group underwent a sham surgery with a midline incision and peritoneal stitches. For the endometriosis group, where we evaluated in vivo LPS^low^ innate immune training (IMM-EDT), mice were administered daily peritoneal injections of low doses (0.1 mg/kg) of LPS for 5 days the week before endometriosis induction. The PBS-EDT group received PBS injection instead. Lesion size was measured in the PBS-EDT and IMM-EDT groups using a high-frequency ultrasound imaging system (Vevo® 2100 VisualSonics; Toronto, CA) at 1 and 3 weeks after surgery.

### Ethical statements

The animal experiment protocol was approved with the ethical approval number DAP20-104 and authorization APAFIS#30327 by the Ministry of Higher Education and Scientific Research.

### Mice pregnancy follow-up

Three weeks after endometriosis induction, CBA/J female mice from the three groups (Sham, PBS-EDT, and IMM-EDT) were mated with DBA/2 male mice. The presence of a vaginal plug was considered embryonic day 0.5 (E0.5). A high-frequency ultrasound imaging system (Vevo® 2100 VisualSonics; Toronto, CA) was used to assess the number of implantation sites (between E7.5 and E10.5) and the number of alive/dead/resorbed fetuses (between E11.5 and E14.5). For ultrasound imaging, pregnant mice were anesthetized with isoflurane and restrained on a heated stage. The fur was removed from the abdomen using hair removal agents, and pre-warmed ultrasound contact gel was applied to the shaved abdomen. The beating heart was detected in the living fetuses. Dead fetuses displayed no beating heart but had visible organ structures. Resorbed embryos/fetuses displayed an echogenic dot with no discernible organ structure. Gestational age was confirmed by observing key developmental features (E8.5 heart, head, and whole embryo; E9.5 amniotic membrane, yolk sac, and cerebral ventricles; E10.5 umbilical cord, placenta, and eyes; E12.5 spine; and E13.5 face, skull bones, and ribs). At the end of gestation (E18.5), the mice were sacrificed to harvest the fetuses, placentas, uterus, and lesion tissues. The number of fetuses, their weights, and the weights of the placentas and endometriosis lesions were recorded. The resorption rate was calculated as (number of implantation sites at the first ultrasound - number of live pups at sacrifice) / number of implantation sites at the first ultrasound × 100. The lesion volume was calculated using measurements obtained by ultrasound (length × width × width).

### Statistical analyses

All data from the mouse experiments were analyzed using GraphPad Prism 8.0 software (GraphPad, San Diego, CA, USA). For the comparison of two groups of data, we used the Student t-test with variables following a normal distribution or the Mann-Whitney *U* test otherwise. Statistical significance was set at p < 0.05.

### HES staining

Endometriosis lesions at E18.5 were retrieved from gestant CBA mice and fixed in 4% paraformaldéhyde (PFA) for 24 hours then PFA was replaced by ethanol 70%. Samples were dehydrated and embedded in paraffin blocks at the histology platform of Institut Cochin in a Logos One (Milestone F/61504/R). Blocks were cut in 5µm sections using a Leica microtome (RM2145).

Slides with the sections were processed in the histology platform for hematoxylin, eosin and saffron staining with a Leica SPECTRA automate.

### RNA sequencing

#### Mouse tissues sampling

The three groups of CBA/J mice were obtained again as described previously and mated with DBA mice three weeks after surgery. Pregnancy was assessed by the presence of a vaginal plug (E0.5 stage), and ultrasound was performed at E7.5-E8.5 to confirm pregnancy. Mice were sacrificed at E9.5, and the feto-maternal interfaces were retrieved from the uterus and either snap-frozen in nitrogen (for snRNA-seq) or digested to isolate immune cells (for CITE-seq). Fetal-maternal interfaces showing signs of abnormal development during ultrasound and/or dissection were not harvested to avoid non-specific signals (considered as dying structures).

#### Nuclei isolation and single nucleus RNA sequencing (snRNAseq)

Nuclei were isolated following a published method^22^. Briefly, materno-fetal interface tissues were lysed for 10 min at 37°C with the lysis buffer described in the method. Samples were dounced with 10 strokes and filtered with a 70µm strainer. The filtrate was centrifuged and filtered again using a 40µm strainer. The filtrate was centrifuged again, and the pellet was resuspended in the staining buffer described in the method with specific antibodies. Nuclei from each placenta were tagged using BioLegend ® TotalSeq ™ antibodies. Nuclei were processed using the Single Cell 3′ Gene Expression kit v3 (10× Chromium, 1000076) according to the manufacturer’s instructions. In brief, 10000-16,000 nuclei per sample were loaded onto Single Cell Chips B to recover as many nuclei as possible (targeting 3,000–10,000 nuclei per sample) while limiting the potential for doublets. Using a Chromium Controller, Gel Bead-In Emulsions were generated, and samples were subsequently processed to isolate and amplify cDNA, and ultimately construct libraries. The quality and concentration of cDNA were evaluated using an Agilent 2100 Bioanalyzer. The quality and concentration of libraries were evaluated by qPCR and on an Agilent 2200 TapeStation, and libraries were sequenced on an Illumina NovaSeq 6000 through the Genom’IC platform of Institut Cochin.

#### Immune cell isolation, surface protein barcoding, single cell RNA and epitope sequencing (CITE-seq)

Previously snap-frozen E9.5 feto-maternal interfaces were digested using a Multi Tissue Tissue Dissociation Kit (Miltenyi Biotec 130-110-201) and a gentleMACS Dissociator (Miltenyi Biotec 130-134-029) at 37°C for 15 min. Samples were filtered using a 100µm strainer and centrifuged to obtain the pellet. The pellet was resuspended in red blood cell lysis buffer (BioLegend # 420301) for 2 min at room temperature. Samples were centrifuged, and the pellet was resuspended in RPMI 2% FCS. Samples were incubated with TruStain FcX PLUS Blocking Reagent (BioLegend 156603) for 10 min at 4°C. Samples were centrifuged, and the pellet was incubated for 30 min at 4°C with a mix of antibodies containing CD45.2 and Zombie NIR (dead/live marker) for sorting, and several epitope markers listed below. Each sample was incubated with a specific antibody tagged with a hashtag (TotalSeq B antimouse), which is listed below. Two wash cycles (centrifugation and addition of RPMI 2% PCF) were performed for each sample. Samples were sorted using a BD FACSAria™ III Cell Sorter keeping all the CD45+ and ZombieNIR-cells. The sorted samples were then processed using the Single Cell 3′ Gene Expression kit v3 (10× Chromium, #1000076) and following the same sequencing protocol as previously described.

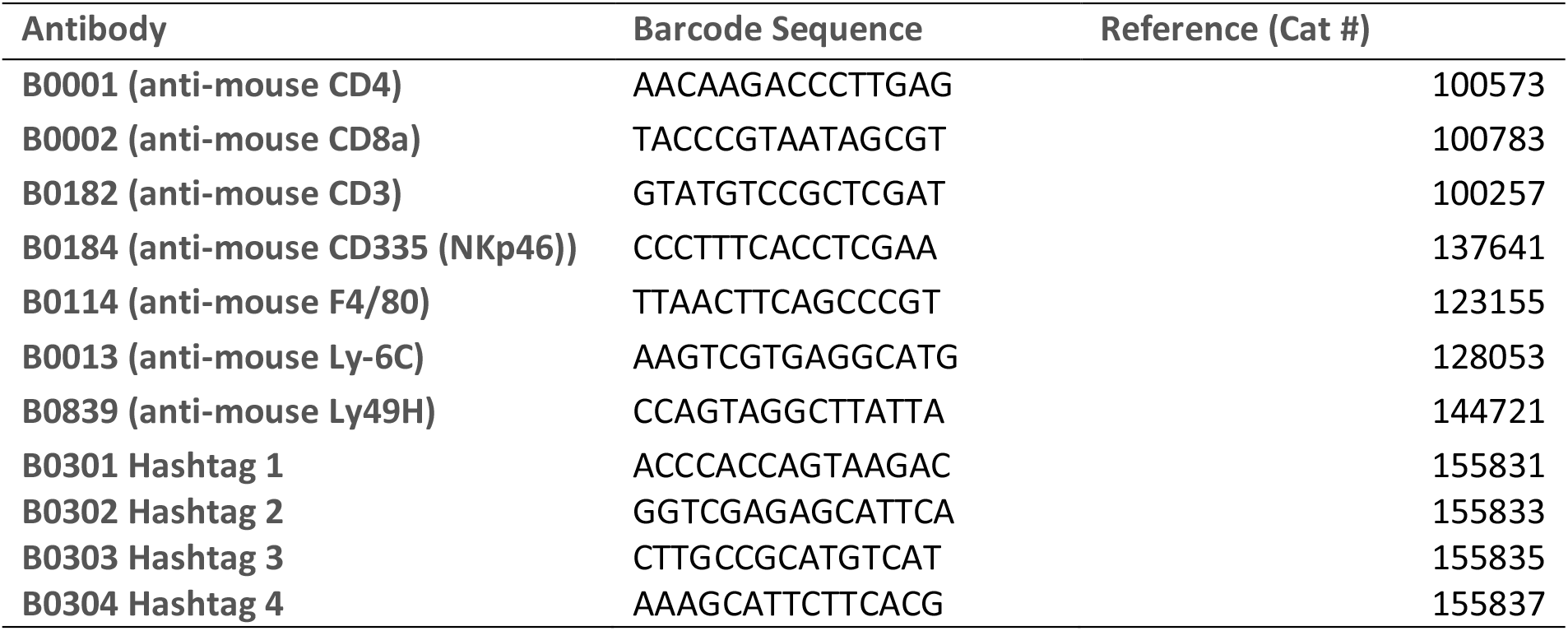

#### Sequences quality control and alignment

Base calling was completed using Illumina® NovaSeq 6000 RTA v3.4.4 software, and BCL base call files were converted to FASTQ files using bcl2fastq conversion software (v2.20). Using the Cell Ranger software suite (v4.0.0) (10X Genomics®), FASTQ files were aligned to the GRCm38 mouse reference genome (modification steps provided by 10X Genomics), and gene-barcode count matrices were generated for all samples.

### Computational analysis of snRNA-seq and CITE seq data

All basic analyses were performed in R using Seurat 4.1.1 package included in a Snakemake pipeline with the following rules (with key parameters described below): quality control, demultiplexing, normalization, integration, and clustering. Muscat package was used to perform differential gene expression analysis. Scripts are available at the bkheira/doridot-sncell GitHub repository.

#### Quality control & demultiplexing

All raw gene-barcode count matrices were converted into Seurat objects in R for initial quality control filtering. Nuclei/cells were filtered using their RNA molecule count (>200 and <30000), sequenced gene count (>500 and < 10000), and percentage of mitochondrial sequenced genes (<10%). Since samples were tagged with antibodies presenting a barcode (TotalSeq anti-mouse hashtags), we could separate them in our analysis by reading these barcodes. An assay “HTO” was created in the Seurat Object and contained barcode data. HTO assay data were normalized and demultiplexed using the “HTOdemux” function of Seurat. Doublet and negative cells for the barcodes were removed from each Seurat object.

#### Normalization & integration

Seurat objects were normalized using SCTransform. To identify the cell populations present in all groups, the Seurat object of each group was integrated with the others. Integration features were selected using the “SelectIntegrationFeatures” function. The SCTransformed data were prepared using the “PrepSCTIntegration” function. Integration anchors were found using the “FindIntegrationAnchors” function, and Seurat objects were integrated using “IntegrateData”.

#### Clustering

Clustering was performed using the “FindNeighbors” and “FindClusters” functions of Seurat with different clustering resolutions. The optimum resolution was chosen using Clustree package in R. Uniform Manifold Approximation and Projection (UMAP) was performed using the RunUMAP function.

#### DEG identification & Gene Set Enrichment Analysis

Differentially expressed genes were identified using Muscat^23^ package in R. Muscat is a package in R that allows pseudobulk analysis by pooling single-cell data in groups of unique clusters per sample. We followed the authors instructions to perform a differential analysis of the mouse data. Data were then filtered : we considered differentially expressed a gene with a p-value adjusted to multiple comparisons smaller than 0.05, an absolute log2FC greater than 0.6 and a minimal expression (logCPM) of 5.

We used the FGSEA package in R using the tutorial available on the Biostatsquid blog^24^. GSEA was performed using the Hallmark database. We added some visualization plots using the package ggplot2.

### Illustrations

Figures 1A and 2A were created using BioRender. BOUZID, K. (2025) https://BioRender.com/trkr2lh

**Figure 1:**
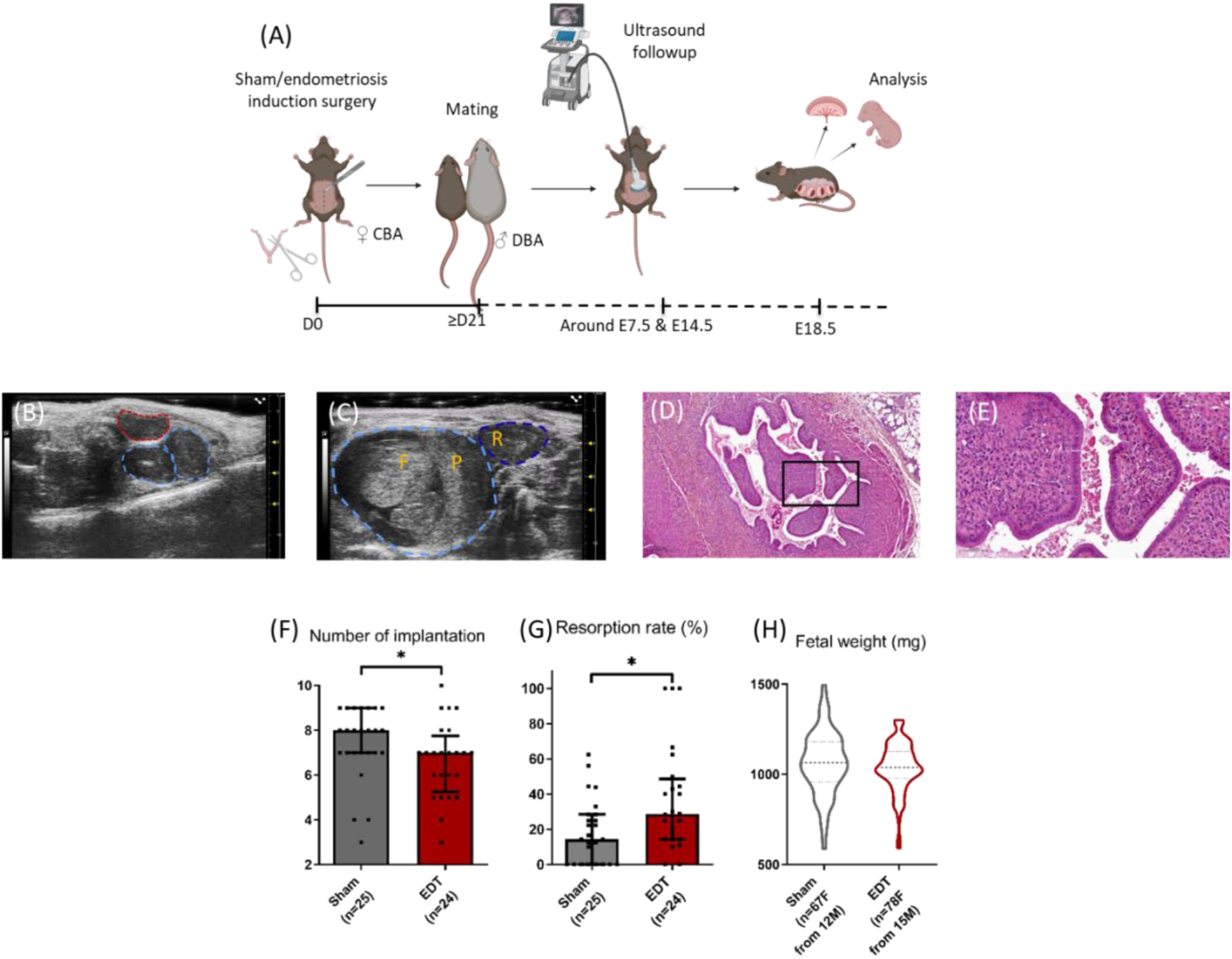
Gestational complications in a mouse model of endometriosis. **(A)** Experimental protocol: briefly, CBA female mice underwent either a sham or an endometriosis (EDT) induction surgery. After 3 weeks of recovery, female mice were mated with DBA males, then gestation and lesion size were followed up by ultrasound at approximately E7.5 and E14.5. At E18.5, one day before parturition, the mice were euthanized, and the fetuses and placentas were retrieved. These are the combined results of three similar experiments. The number of mice is indicated in the figure by n, with a precision of fetuses (F) and gestant mice (M), if relevant. **(B)** Ultrasound image showing two embryos at E7.5 (blue) and an endometriosis like lesion (red) **(C)** Ultrasound image showing alive (light blue) fetus (F) and placenta (P) and resorbed fetus (R, dark blue) at E14.5 **(D)** and Micrographs of Hematoxylin-Eosin-Saffron staining of histological sections from retrieved endometriosis lesions **(F)** Number of implantations per mouse **(G)** Fetal resorption rate per mouse. For and **(G)**, data are presented as median ± interquartile range. **(H)** Fetal weight per pup at E18.5 v*: p-value < 0.05 (Mann-Whitney *U* test).

**Figure 2:**
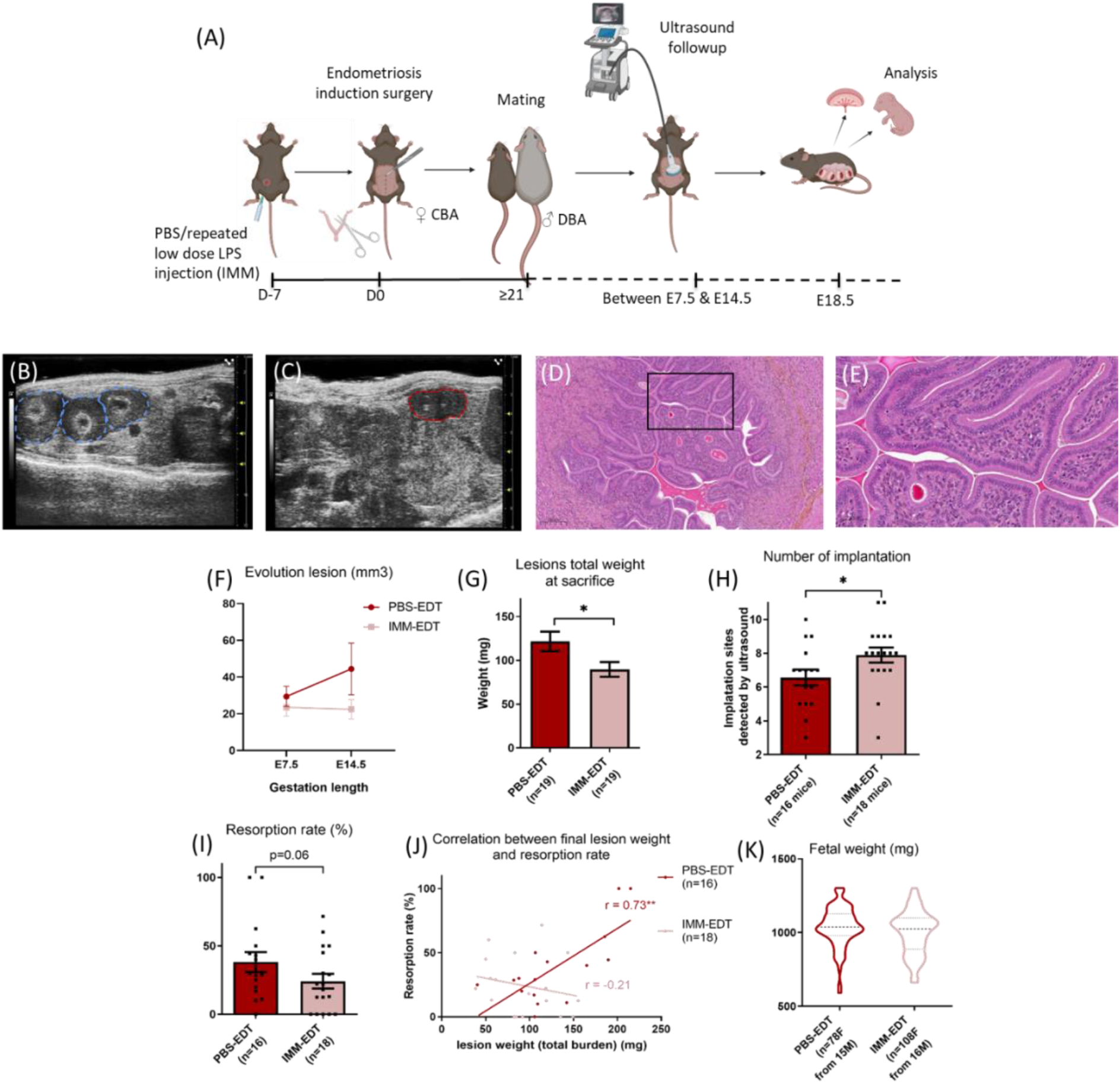
Immunomodulation (IMM) treatment partially corrects gestational complications in endometriosis mice. **(A)** Experimental protocol: repeated injections of PBS or low-dose LPS were administered before inducing endometriosis by surgery, followed by the same protocol as shown in Figure 1A. **(B)** Ultrasound image showing three embryos at E8.5 (blue) and **(C)** an endometriosis like lesion (red) **(D)** and **(E)** Micrographs of Hematoxylin-Eosin-Saffron staining of histological sections from retrieved endometriosis lesions **(F)** Evolution of the lesion volume as observed on ultrasound. For **(G), (H)** and **(I)**, data are presented as mean ± SEM. PBS-EDT: group receiving PBS and two lesions of endometriosis. IMM-EDT: group receiving LPS at low doses and with two lesions of endometriosis. **(G)** Total weight of the two lesions at the time of sacrifice. Student’s t-test **(H)** Number of implantations per mouse, Student’s t-test **(I)** Fetal resorption per mouse, Student’s t-test **(J)** Correlation between total lesion weight and resorption rate per mouse, PBS-EDT Pearson r = 0.72 (p = 0.0013), LPS-EDT Pearson r = -0.21 (p = 0.38). Fetal weights. *: P-value < 0.05, **: P-value < 0.01.

## Results

### Endometriosis reduces the number of embryo implantations and increases fetal resorption rate in the CBA/JxDBA/2 mouse model

We first wanted to check if the presence of endometriosis-like lesions had a deleterious effect on pregnancy in the CBA/JxDBA/2 mouse model, as it has been demonstrated in other mouse strains in the context of syngenic crosses (C57BL/6^9,10^, Swiss Webster^11^ or Balb/C^12^). Our experimental design (**Figure 1A**) was as follows: we induced endometriosis in CBA females by surgically suturing two syngeneic uterine fragments on the peritoneal wall, as previously described^15^. This model allows us to follow the lesions using ultrasound. Three weeks later, the mice were mated with DBA males. Ultrasound imaging was performed (**Figure 1B**) to count the number of implantation sites (between E6.5 and E8.5) and evaluate pup development or resorption at each site (**Figure 1C**, between E13.5 and E15.5). The volume of the endometriosis lesions was also measured. At E18.5, the mice were sacrificed, and the number of live fetuses, their weight, and the weight of the endometriosis lesions were recorded.

Endometriosis lesions had a typical endometrium structure with an monolayered epithelium and a dense stroma (**Figure 1D and 1E**). Mice with endometriosis lesions displayed a slight but significant decrease in the number of implantations (10% decrease, p=0.03, **Figure 1F**). A 2-fold increase in the resorption rate in the group with endometriosis lesions (18.9 % vs. 36.9%, p=0.047, **Figure 1G**) was also observed. This effect seems to be dose-dependent, as mice with only one endometriosis lesion showed an intermediate non-significant increase in resorption rate (fold change of 1.5 with 27.8% resorption rate, p=0.14, **Supplementary Figure 1**). Fetal weights at E18.5 were equivalent in all groups (**Figure 1H**, Mann-Whitney test, p = 0.32). Histological evaluation of E18.5 placentas did not reveal any obvious anomalies associated with endometriosis (**Supplementary Figure 2**).

### Pre-conceptional introduction of low-dose LPS reduces gestational complications associated with endometriosis in a mouse model

As we previously showed that immunomodulatory (IMM by repeated exposure to low doses of LPS) treatment can reduce the lesion size in endometriosis mice^15^, we wanted to see if this immunomodulatory treatment could also reduce the deleterious effect of endometriosis on pregnancy. We conducted an experiment (**Figure 2A**) in which endometriosis induction was preceded by peritoneal injections of repeated low doses of LPS (IMM) or PBS. Three weeks after surgery, the endometriosis mice were mated with DBA/2 males, and gestation was followed up as previously described. We previously showed that IMM has no effect on the resorption rate in mice without endometriosis^21^.

The volume of the lesions was measured during ultrasounds following gestation, and IMM treatment showed a trend toward reducing the volume of the lesions (**Figure 2F**, p=0.15 by mixed effect analysis). At the end of gestation, the lesion weight of the IMM-EDT group was significantly lower than that of the PBS-EDT control group (**Figure 2G**, p=0.01). Lesions of IMM-EDT group had endometrial glands structures (**Figure 2D and 2E**), similar to those observed in **Figure 1D**. The number of implantations was significantly higher in the IMM-treated group (20% increase, **Figure 2H**, p = 0.048). The fetal resorption rate was 38% in the PBS-treated group and 24% in the IMM-treated group **(Figure 2I**, p = 0.06), which is closer to that observed in mice without endometriosis (sham group in **Figure 1E**, 19%). The significant correlation between lesion size and resorption rate in the endometriosis group (r=0,73, p=0.0013) was lost with the IMM (**Figure 2J**). Fetal weights at E18.5 were equivalent in all groups (**Figure 2K**).

### SnRNAseq reveals an increase of Gata4 and a decrease of Prap1 expression in decidual cells in endometriotic mice

Since there are a decreased mean number of implantation sites and an increased resorption rate in the endometriosis group, we hypothesized that there is a possible dysfunction of the decidua and/or the placenta that leads to an altered or interrupted embryo development. To validate this hypothesis, feto-maternal interfaces at E9.5 without visible signs of ongoing resorption were collected from mice with sham surgery or with two lesions of endometriosis, and their transcriptomes were sequenced at the single-nuclei level.

We identified 13 cell clusters in our dataset that were distributed homogeneously among the samples (**Figure 3A, 3B**). *Wt1*, a known marker of decidualized cells^25^, was found in seven clusters (**Figure 3D**). Among them, there are two decidual fibroblast/stromal clusters, identified by the expression of collagen genes (*Col3a1*) and several genes involved in the remodelling of the extracellular matrix of the decidua during implantation and placentation (*Sulf1, Adamtsl1*)^26,27,28^. Interestingly, these decidual fibroblasts expressed *Il15* and its receptor *Il15ra*, which are involved in the activation of uterine Natural Killer (uNK)^29,30^. The other *Wt1*+ clusters were identified as decidual stromal cells (DSC) as they expressed several known markers of decidual stromal cells at different levels (*Hand2, Pgr*, and *Esr1*), suggesting that the clusters may represent different stages of differentiation or different subtypes of decidual stromal cells. All clusters’ markers are available in **Supplementary Table 1**.

**Figure 3:**
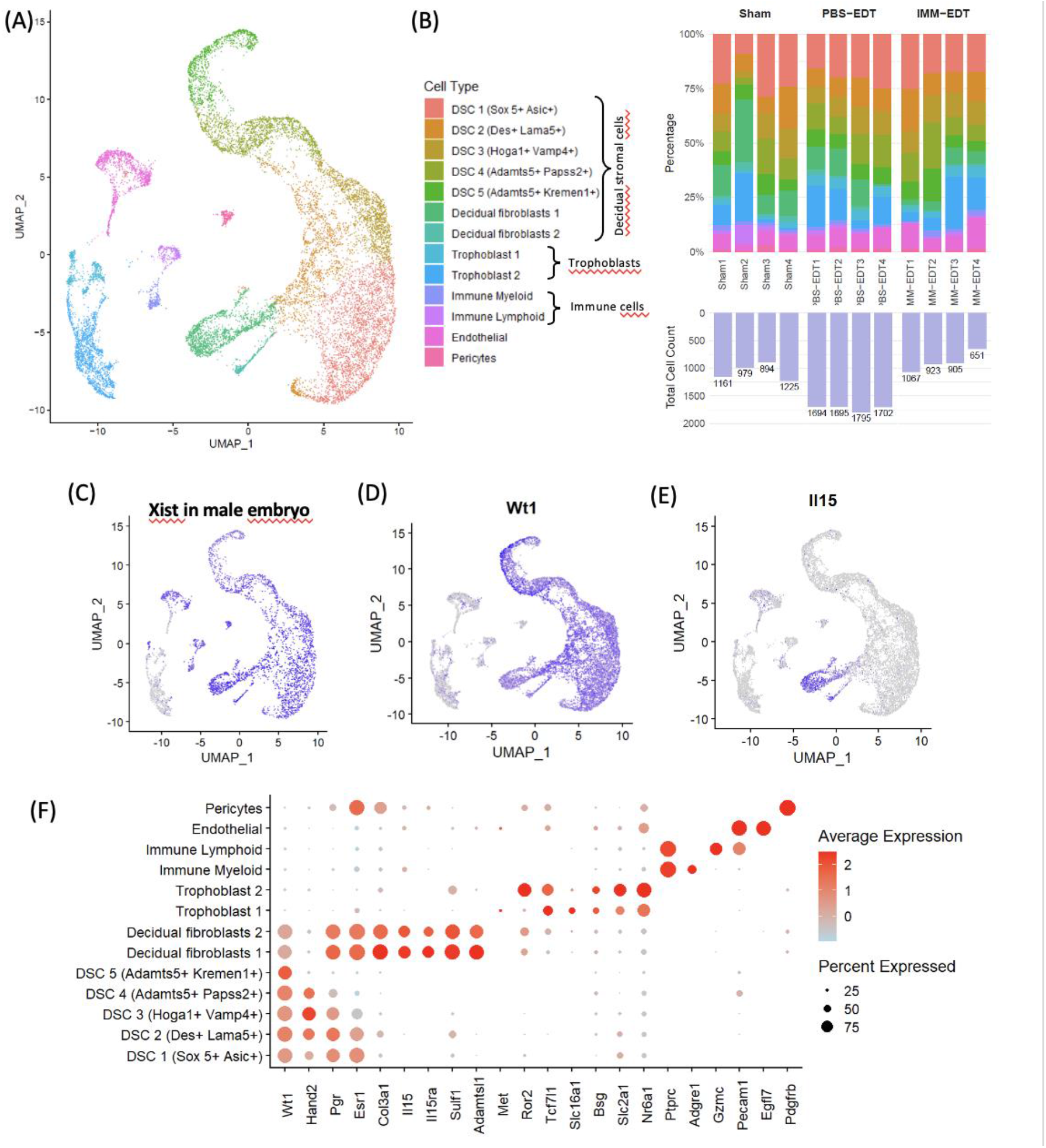
Single-nucleus RNA sequencing of E9.5 fetomaternal interface reveals its composition and characteristics. (A) UMAP of cell transcriptomes (B) Graph representing the proportion of a cell type (cluster) in each sample (top) and the number of cells retained for analyses after quality control filtering (bottom) (C) Expression of *Xist* gene in materno-fetal interfaces with male embryos (D) Expression of *Wt1* (E) Expression of *Ptprc* (CD45 gene) (F) Dotplot of cluster identifying genes. DSC: Decidual stromal cells

The expression of the maternal gene *Xist* in placental samples associated with a male embryo segregated the cells of embryonic origin from those of maternal origin (**Figure 3C**). We confirmed their trophoblast origin by the expression of specific genes that mark them as trophoblast progenitors, such as *Epcam* and *Ror2* expression in embryonic cells^31, 32^ and trophoblast cells with *Mct1/Sl16a1, CD147/Bsg, Glut1/Slc2a1*^33^, and *Nr6a1*^34^ (**Figure 3F**). The expression of *Pecam1* was used to identify endothelial cells within the placenta^35,36^. Cells of hematopoietic origin were identified by *Ptprc* (CD45 gene, **Figure 3E**). Pericytes have been identified using the Pdgfrb gene.

Through differential expression analysis (**Figure 4**), we identified several differentially expressed genes in all clusters. When plotting a heatmap of the top 10 DEG in each cell group, we observed that the transcriptomic expression of the IMM-EDT group was often between the expression of the control and endometriosis groups, indicating a partial correction of the gene expression differences induced by endometriosis by IMM treatment. In the different clusters of decidual stromal cells, we observed two genes that were consistently differentially expressed: *Gata4* was upregulated and *Prap1* was downregulated in the group with endometriosis-like lesions. Interestingly, these 2 genes differential expression were normalized by the immunomodulatory treatment, with a decrease of *Gata4* and an increase in *Prap1* in the IMM-EDT group compared to the PBS-EDT group. A list of all differentially expressed genes is available in **Supplementary Tables 2 and 3**.

**Figure 4:**
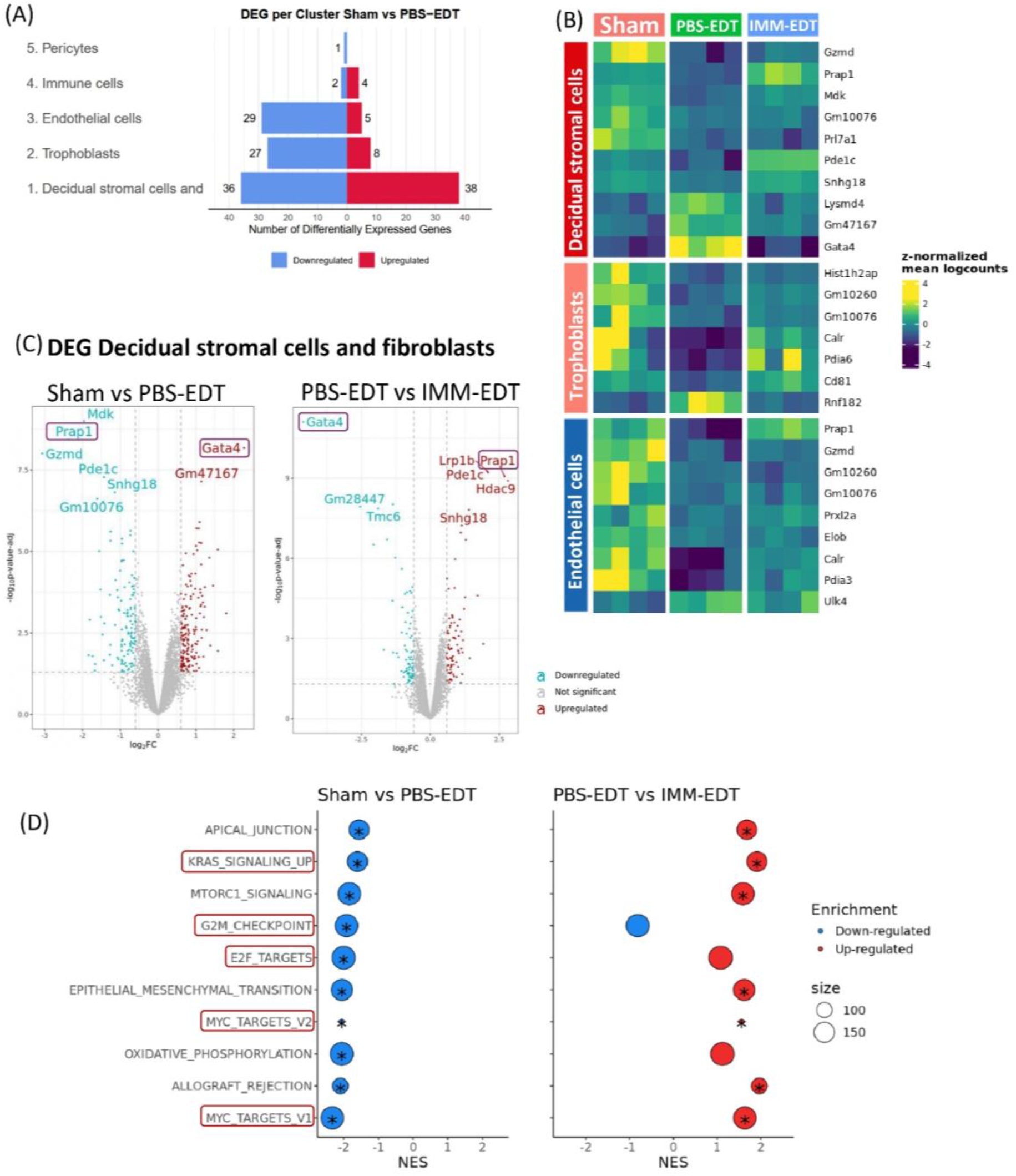
Differential expression analysis reveals alterations driven by endometriosis and partially corrected by the IMM treatment. (A) Number of differentially expressed genes per group of cells (B) Heatmap of the top 10 differentially expressed genes between the sham and endometriosis groups. Cell clusters were grouped by cell type as shown in Figure 3B. (C) Volcano plot showing the differentially expressed genes within the decidual stromal cells (D) Dotplot of Gene Set Enrichment Analysis: top 10 significantly enriched pathways from Sham vs PBS-EDT analysis and how they are in the PBS-EDT vs IMM-EDT in the pooled DSC group. Pathways involved in proliferation are surrounded by red rectangles. * Significant enrichment after adjustment.

For the next analysis, we grouped the clusters by cell type (decidual stromal cells, trophoblast cells, endothelial cells, immune cells, and pericytes) and compared the groups of mice (sham, PBS-EDT, IMM-EDT) by performing differential expression analysis and gene set enrichment analysis (GSEA). We see transcriptomic changes induced by endometriosis in all the cell types (**Figure 4A, B**). We then chose to focus on decidual stromal cells, which are the group of cells with the most differentially expressed genes induced by the presence of endometriotic lesions (**Figure 4A**). As expected, the differential expression analysis showed an overexpression of Gata4 (log2FC of 2.27 and adjusted p-value of× 10^−5^) and a downregulation of *Prap1* (log2FC of -2.37 and adjusted p-value of 1.63 × 10^−5^1.63) in the PBS-EDT group compared to the Sham group (**Figure 4C**). IMM led to the correction of these expressions (**Figure 4B**). GSEA showed that pathways related to proliferation (Myc targets, E2F targets, G2M checkpoints, and KRAS signalling) were negatively enriched in the endometriosis group. This downregulation was significantly corrected by IMM treatment, except for the G2M checkpoint (**Figure 4D**).

### CITEseq analysis reveals transcriptomic differences in all immune cell types and an inflammation in endometriotic placentas

After immunomodulation, it is relevant to examine the consequences on immune cells directly. As these cells account for a small proportion of the maternal fetal interface and thus are not well represented in the whole interface snRNAseq dataset, we performed Cellular Indexing of Transcriptomes and Epitopes by Sequencing (CITEseq or proteotranscriptomics) on isolated immune cells (CD45+) in the E9.5 materno-fetal interface.

We identified 11 clusters of cells in this dataset (**Figure 5A)**, and the cell types were equally represented in the samples (**Figure 5B**). A major group of myeloid cells was recognized by *Cd68* and *Itgam* (CD11b) expression (**Figure 5G**) and the expression of the monocyte marker Ly6C (**Figure 5D**). Within this group, we identified dendritic cells expressing *Cd209a* and a small cluster of plasmacytoid dendritic cells expressing *Siglech*. Neutrophils were identified by the expression of calprotein genes (*S100a9* and *S100a8*). Macrophages, identified here with the antibody F4/80 (**Figure 5C**), had several marker genes, such as *Wwp1, Ccl8, Stab1, Gas6*, and *Fcrls*. Three other clusters were identified as monocytes (all expressing Ly6C and/or *Itgam*), and specific genes differentiating these subcategories of monocytes were identified.

**Figure 5:**
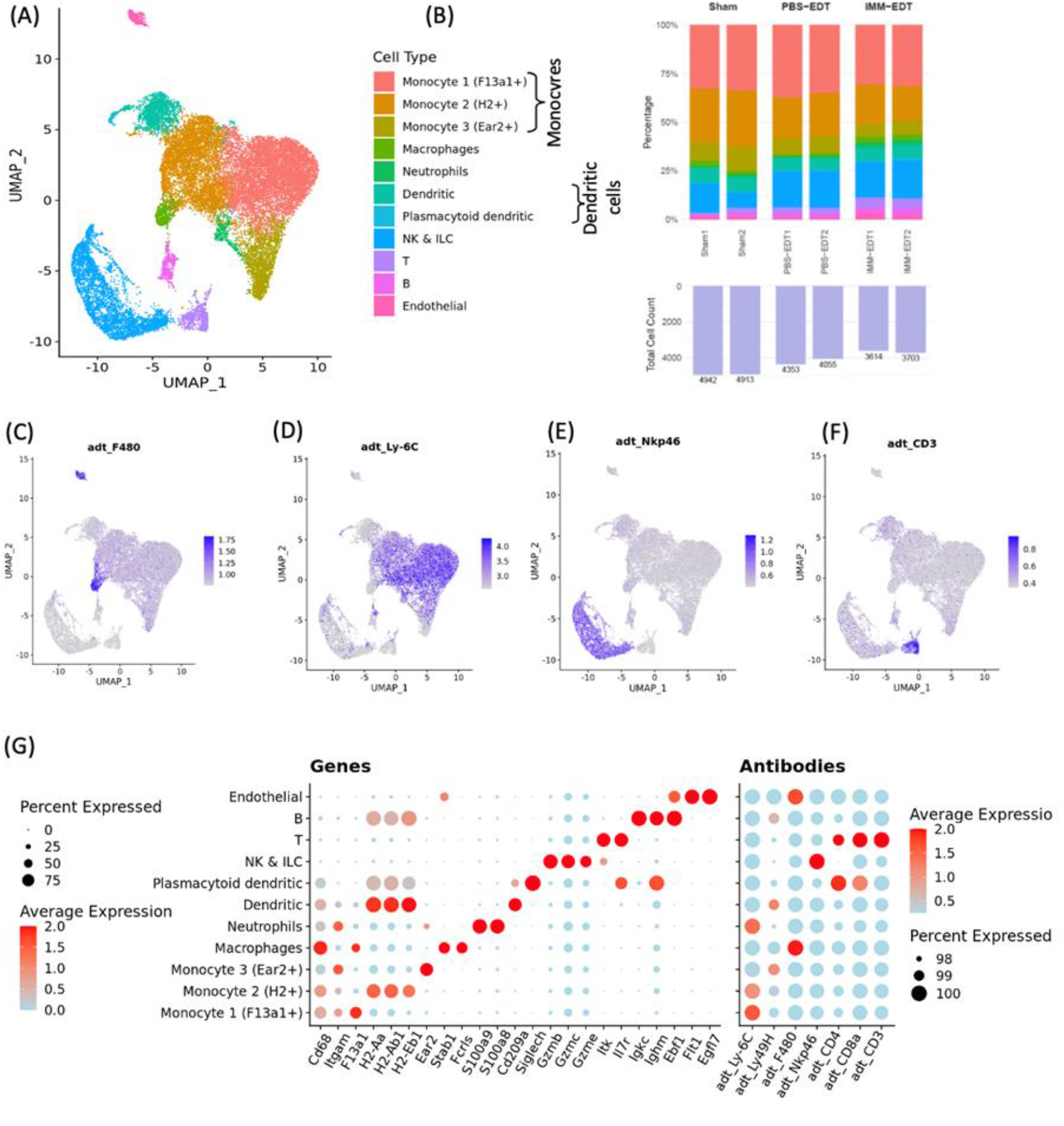
Composition of E9.5 fetomaternal interface by single cell proteotranscriptomics sequencing of immune cells. (A) UMAP of cell transcriptomes (B) Graph representing the proportion of a cell type (cluster) in each sample (top) and the number of cells retained for analyses after quality control filtering (bottom) (C) Featureplot of the surface antibody F4/80 staining macrophages (D) Ly-6C (E) Nkp46 (F) CD3 (G) Dotplot of cluster identifying genes and antibodies

The other major cell type was natural killer (NK) cells and innate lymphoid cells (ILC) identified by the different granzyme genes (*Gzmb, Gzmc, Gzme*) and perforin (*Prf1*) and with the surface protein data of the CITEseq, using the NKp46 antibody (**Figure 5E**). T cells were identified by the surface proteins CD3 (figure 5F), CD4, and CD8a, and at the transcriptomic level, they had specific expressions of *Itk* and *Il7r*. B cells were identified with immunoglobulin genes such as Igkc or Ighm, as well as genes like *Ebf1, Bank1, Cd79a*, and *Cd79b*.

Finally, in the CD45+ cells, we also had a small cluster of endothelial cells recognized by the expression of known genes such as *Flt1* and *Egfl7*. Cluster markers are available in **Supplementary Table 5**.

We could see in the PBS-EDT group compared to the Sham group few transcriptomic differences in all the cell types (**Figure 6A**). The IMM treatment induced a higher number of significant transcriptomic differences in all cell types compared to the Sham group (**Supplementary Figure 3A**) and to the PBS-EDT group (**Figure 6A**). Indeed, immunomodulation of the immune system leads to transcriptional changes^37^. A gene that was consistently downregulated among the cell groups (except macrophages) was *Slc15a2* (**Figure 6B**), which encodes the peptide transporter PEPT2 (see **Supplementary Table 7**).

**Figure 6:**
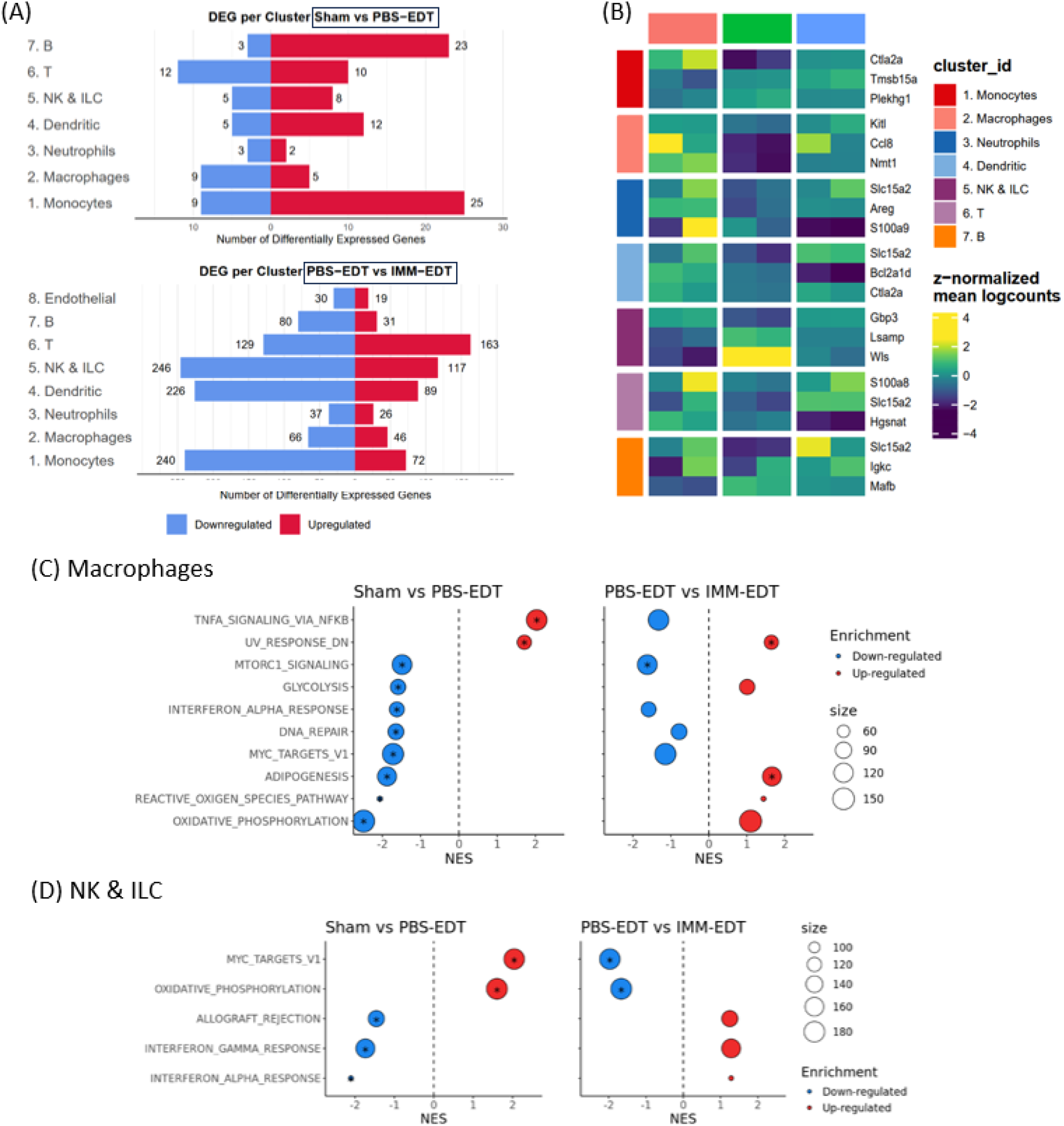
Differential expression analysis in immune cells reveals alterations driven by endometriosis and partially corrected by the IMM treatment. (A) Number of differentially expressed genes in each cell group. (B) Heatmap of the top 3 differentially expressed genes between the sham and endometriosis groups. Cell clusters were grouped by cell types as shown in Figure 5A. (C) Dotplot of Gene Set Enrichment Analysis : top 10 significantly enriched pathways from Sham vs PBS-EDT analysis and how they are in the PBS-EDT vs IMM-EDT in Macrophages and (D) in NK and ILC. * Significant enrichment after adjustment.

To determine the pathways in which these genes are involved, we performed GSEA. Notably, we observed an upregulation of TNFα signalling in macrophages in endometriosis, which tended to be corrected by IMM treatment (**Figure 6C**). This upregulation is also observed in endometriosis in the pooled group of monocytes (**Supplementary figure 3**), in T and B cells, but only corrected by the IMM in B cells (see **Supplementary Table 8**). In NK and ILC, endometriosis led to a negative enrichment of IFNγ and IFNα response pathways, and IMM tended to normalize this pathway (**Figure 6D**). This negative enrichment of IFNγ response in endometriosis is also found in decidual stromal cells, endothelial cells, and trophoblasts.

## Discussion

In this study, we first examined the impact of endometriosis-like lesions on gestation in CBA/JxDBA/2N crossing. Endometriosis-like lesions in our mouse model have structures similar to those found in humain endometriosis^38^, with a glandular aspect. Endometriosis led to fewer implantations and a higher proportion of fetal resorptions than in the sham group. Other studies have shown the consequences of endometriosis on gestation in mouse models, including a reduction in the pregnancy rate^9,10,12^ (gestant mice per group), increase in the resorption rate^11,12^, decrease in litter size^12^, and reduced birth weight for the pups^11^. In our study, we found no significant difference in fetal weight, which may be due to the fact that mice were sacrificed before parturition; however, other studies also showed no difference in pup weight at E18.5^10^. Currently, most of these studies on mouse models have been used as preclinical models to test the effects of potentially beneficial molecules, but the underlying mechanisms of these complications are yet to be understood. Our study suggests that the presence of lesions and/or inflammation induced by endometriosis leads to a deleterious environment for pregnancy development.

This is further substantiated by the improvement in pregnancy outcomes with IMM treatment. In accordance with previous findings of our team, inducing immunotolerance reduced the evolution and final size of endometriosis lesions and had a beneficial effect on pregnancy complications, with an increase in the number of implantations and a decrease in the fetal resorption rate. In the IMM condition, the correlation between lesion size and fetal resorption was lost, suggesting that the beneficial effect of IMM on pregnancy may be due to a reduction in inflammation rather than a reduction in lesion size.

Due to the increased risk of miscarriage in endometriosis, there may be defects in early pregnancy. To study this, we used our mouse model again and retrieved the feto-maternal interface at E9.5, a stage at which the structure of the placenta is developing. Transcriptomic sequencing has shown that endometriosis induces changes in all cell types of the developing placenta, even in trophoblasts, which are exclusively fetal. These fetal cells are derived from oocytes exposed to chronic inflammation induced by endometriosis, which may explain the consequences on trophoblastic cells. These changes in fetal cells of the placenta raise questions about the transmission of endometriosis consequences to the fetus via epigenetic signals. Indeed, endometriosis has a genetic heritability of 50%, and until now, several GWAS have only been able to explain 5%^39^ of the genetics of endometriosis. In addition to potential rare variants not detected by GWAS, missing heritability could also be due to epigenetics and should be studied in further research. Our model may allow for the exploration of its potential impact on the next generation.

The major cell type at E9.5 in the developing placenta is decidual stromal cells (DSC), which are derived from the decidualization of endometrial stromal cells. We observed most transcriptomic changes in the presence of endometriosis in this group of cells. *Gata4* gene expression was upregulated by endometriosis and normalized by IMM treatment. This upregulation of *Gata4* has already been found in endometriosis within ectopic and eutopic endometrium in patients^40^, making our mouse model relevant for studying this upregulation. The involvement of *Gata4* in gestation has not been studied, but its family members GATA2 and GATA3 are important during preimplantation and early post-implantation development, with knockouts of both *Gata2* and *Gata3* leading to placental defects^41^ in trophoblasts.

DSC also showed downregulation of Prap1 in endometriosis, which was corrected by IMM treatment. In mice, Prap1 appears to be involved in establishing uterine receptivity to the embryo^42^ and is an indicator of successful implantation^43^. Therefore, its downregulation may participate in the reduced number of implantations and increased resorption observed in endometriotic mice. Prap1 is a gene positively regulated by the progesterone receptor PGR in the mouse endometrium^44^. *Pgr* expression in our dataset was non-significantly downregulated by endometriosis (logFC of -0.40, p-value of 0.0044, and adjusted p-value of 0.077) and may participate in the downregulation of *Prap1* expression. This downregulation of *Pgr*, although not significant in our dataset when we adjusted the p-value for multiple testing, should be further studied, and our model may be interesting to study the progesterone resistance found in endometriosis in human^45^.

In our GSEA of mouse DSCs at E9.5, most of the downregulated pathways in the endometriosis group were related to proliferation. A defect in proliferation during early gestation could explain miscarriages or further placental pathologies found to be increased in endometriosis. We identified two clusters of decidual stromal cells that expressed extracellular matrix remodelling genes and named them decidual fibroblasts. These decidual fibroblasts expressed *Il15*, which encodes IL-15, an important factor for the differentiation and proliferation of uNK cells, allowing uNK cells to support embryo implantation, spiral artery remodelling, and immune tolerance during pregnancy^46^. Until now, it was known that IL-15 signalling was coming from decidual stromal cells^30^, but here we show that only a specific subtype of DSC has this role. Interestingly, these cells also express *Il15ra*, a gene encoding a receptor for IL-15, suggesting the possibility of an autocrine signal within these cells.

When sorting the immune cells from the E9.5 feto-maternal interfaces, we identified a majority of myeloid cells and up to 20-25% of lymphoid cells, concordant with previous littereature^47^. This is different from human early placentas, where NK cells represent 70% of immune cells^48^ and macrophages represent 20 and 30%^49^. Previous studies have shown that less than 1% of immune cells are of fetal origin at E10.5^47^, a later stage than that in our study; therefore, it is likely that the vast majority of cells in our dataset are of maternal origin.

Endometriosis induced transcriptomic changes in all immune cells. When focusing on macrophages in endometriosis, GSEA showed an upregulation of an inflammatory pathway, TNFα via NFκB. In endometriosis, it has been shown that macrophages are more activated and participate in inflammation within the peritoneal cavity^50^. Our results suggest that macrophages are inflammatory, even at the maternal-fetal interface. We can imagine that inflammation at an early stage of gestation could lead to a poor establishment of the maternal-fetal interface and thus induce miscarriages/resorptions. The upregulation of inflammatory pathways was no longer observed when endometriosis mice were previously trained with low doses of LPS (IMM). We have already shown that peritoneal macrophages in a mouse model of endometriosis become more immunotolerant with LPS training at low doses^15^, and we show here that this immunotolerance is also induced in the maternal-fetal interface.

The other major immune cells during pregnancy are NK cells. In these cells in endometriosis mice, the IFNγ and IFNα response pathways were downregulated, whereas the Myc target V1 pathway, which is involved in proliferation, was upregulated. uNK cells are the main source of IFNγ in the murine materno-fetal interface^51^. IFNγ is essential for the development of gestation in mice and participates in NK maturation^52^. Thus, NK cells, which may be less able to respond to this signal, may not mature well and may disrupt the development of pregnancy. Mice that are IFNγ^-/-^ present a high rate of fetal resorptions, no spiral artery remodelling, and decidua necrosis. They also have more uNK at the MFI, as if they are trying to compensate for the lack of IFNγ by having more producing cells^51^. The upregulation of a proliferation pathway in NK cells in our dataset may be a consequence of the downregulated IFNγ response. IFN-α and IFN-β are regulators of NK cell activation and their production of IFNγ^53^, and in mice that are *Ifna*-/-, the same phenotype as I*fng*-/-is observed at the materno-fetal interface, suggesting that NK cell IFN-α response is also essential for the development of the MFI. This downregulated response to IFNγ was also found in the decidual cells, endothelial cells, and trophoblasts of our dataset, and only the downregulation of IFNα was found in decidual cells and trophoblasts. Given the importance of IFNγ signaling in vascular remodelling and maintenance of the decidua, this general downregulation of the response may be a cause of disruption of gestational development.

Of note, B cells are among the immune cells with most differentially expressed genes in endometriosis compared to sham group. GSEA showed enrichment of several pathways involved in inflammation and immune response. B cells involvement in endometriosis is still incompletely defined and controversial with evidence supporting increased B cell activation and auto-antibody production^54^. Our mouse model could be used to further study B cells in endometriosis.

Our results show that a murine model is pertinent for studying gestational complications induced by endometriosis and reveals mechanisms affecting decidual stromal and immune cells at the beginning of gestation. Our findings suggest that targeting the immune system may be a therapeutic strategy to improve pregnancy outcomes and reduce the risk of miscarriage.

## Acknowledgments and funding

We would like to thank the different platforms at the Institut Cochin and Université Paris Cité that helped us generate and analyze our data: GENOM’IC, CYBIO, HIST’IM, BIOINFORMAT’IC, and IPOP-UP.

We would like to thank Université Paris Cité for the IDEX funding of the PlacentAtlas project (ANR-18-IDEX-0001).

K. B. received a PhD fellowship from BioSPC doctoral School at Université Paris Cité. C. M. is supported by ANR-20-CE14-0004. L.D. is funded by the European Union (European Research Council (ERC) starting grant No 101078556). Views and opinions expressed are however those of the author(s) only and do not necessarily reflect those of the European Union. Neither the European Union nor the granting authority can be held responsible for them.

