## Supplementary material for "Preconceptional immunomodulation partially corrects pregnancy abnormalities induced by endometriosis in a mouse model, with normalization of transcriptional alterations observed in the developing fetal-maternal interface at the single cell level"

#### Supplementary Material and methods

##### Histological stainings

Feto-maternal interfaces at E9.5, placentas and endometriosis lesions at E18.5 were retrieved from gestant CBA mice and fixed in 4% paraformaldéhyde (PFA) for 24 hours then PFA was replaced by ethanol 70%. Samples were dehydrated and embedded in paraffin blocks at the histology platform of Institut Cochin in a Logos One (Milestone F/61504/R). Blocks were cut in 5µm sections using a Leica microtome (RM2145). Slides with the sections were processed in the histology platform for hematoxylin and eosin or hematoxylin, eosin and saffron staining with a Leica SPECTRA automate.

### Supplementary figures

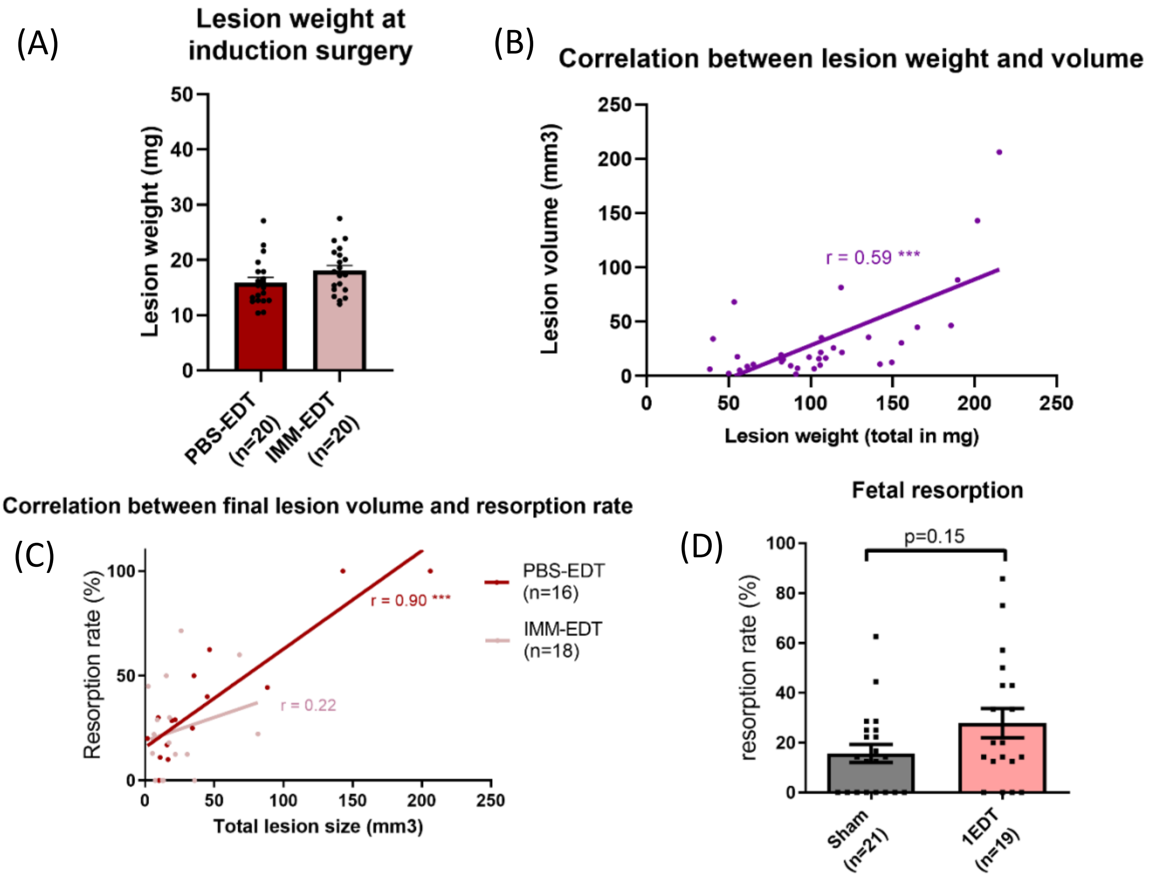


**Supplementary Figure 1: Mouse model of endometriosis**

(A) Weight of the uterine horn pieces before surgery (B) Correlation between the lesion weight sacrifice and lesion volume seen by ultrasound 3 days before sacrifice. Spearman r showed on the graph (C) Correlation between lesion volume and resorption rate. Spearman r showed on the graph. (D) Fetal resorptions depending on the number of endometriosis lesion, 1EDT having one lesion and PBS-EDT having 2 lesions. Kruskall-Wallis test. *: p-value < 0.05, ***: p-value < 0.001

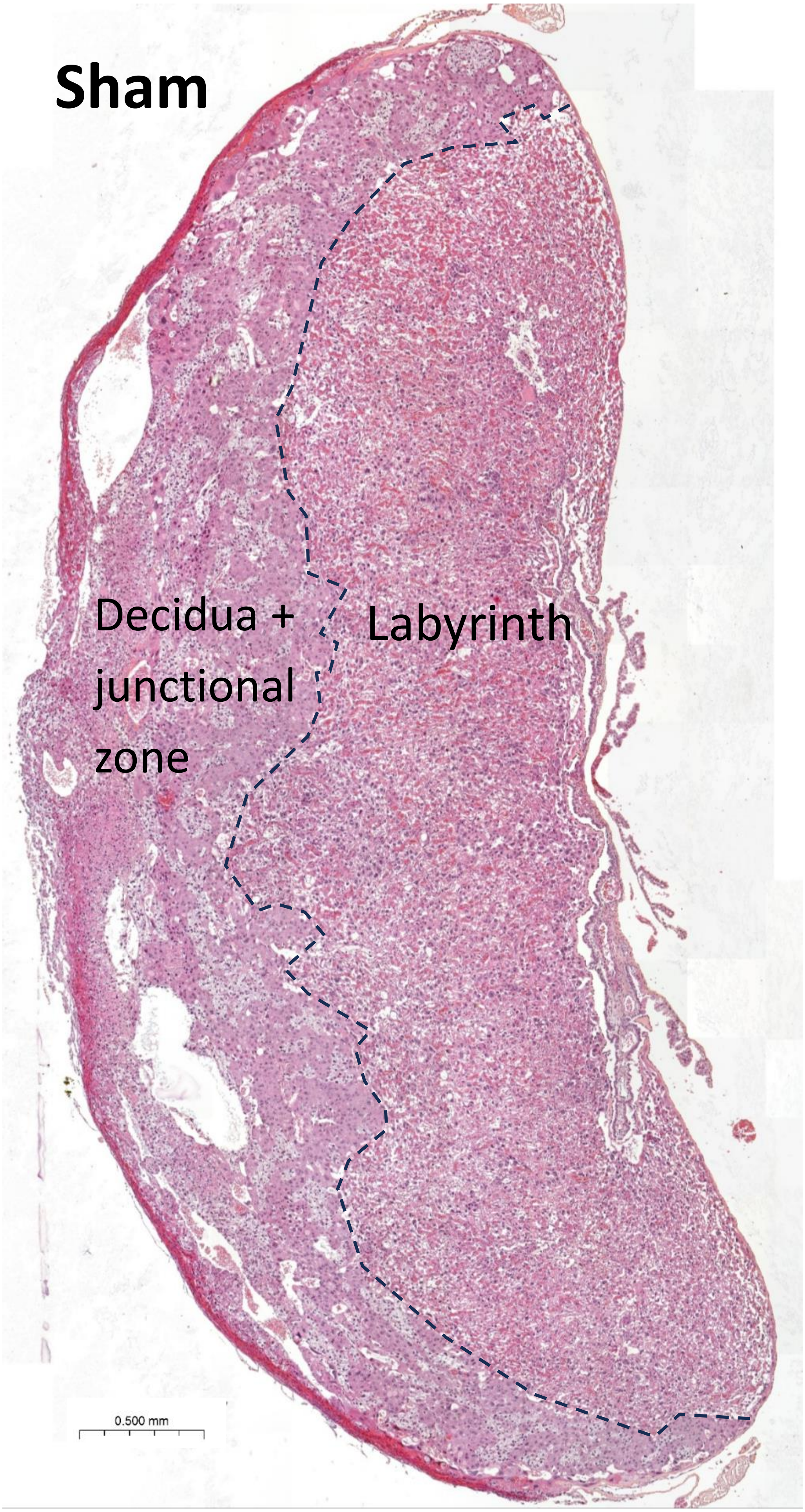
Sham
Decidua +
junctional
zone
Labyrinth
0.500 mm

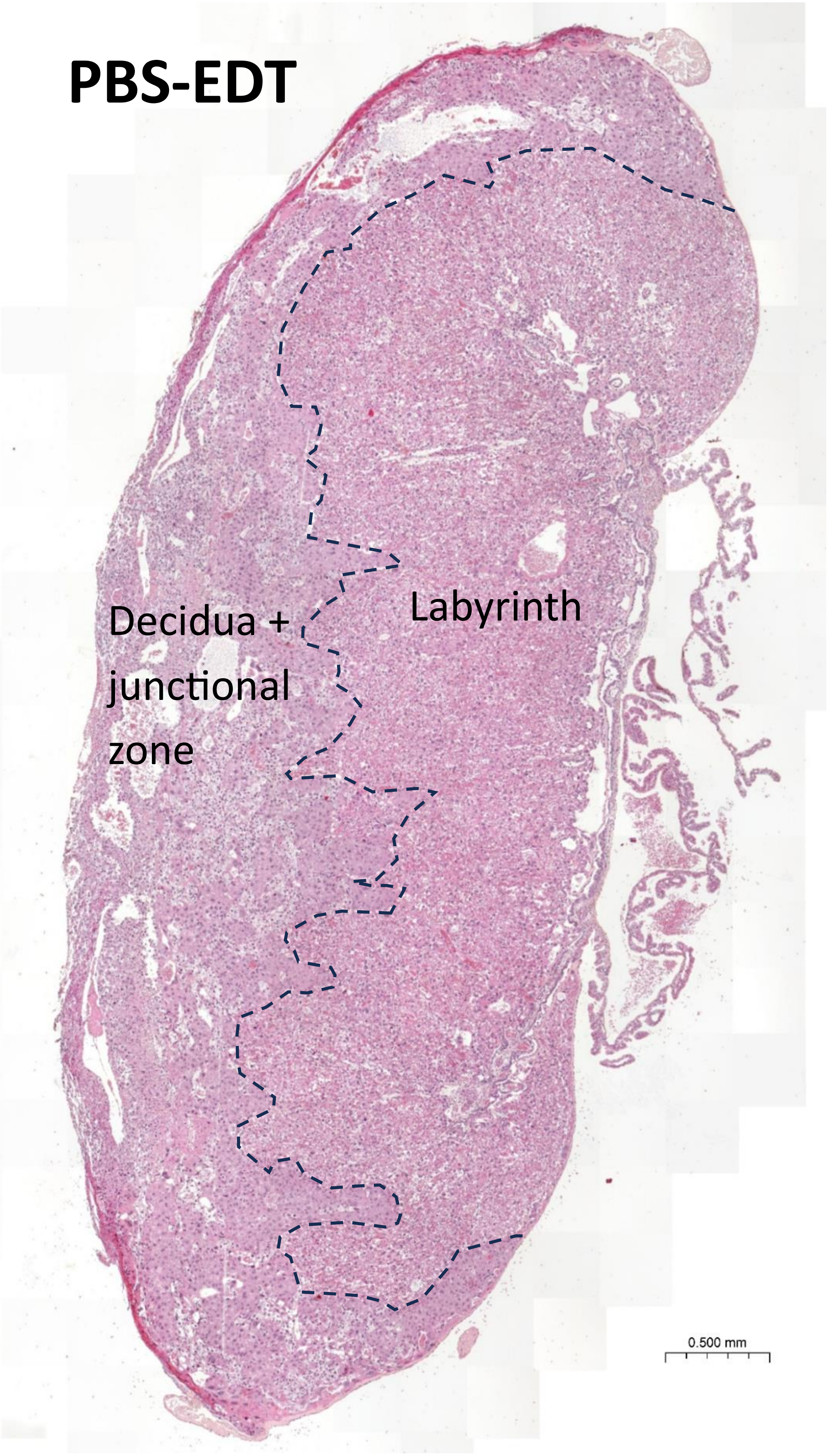
PBS-EDT
Decidua + junctional zone
Labyrinth
0.500 mm

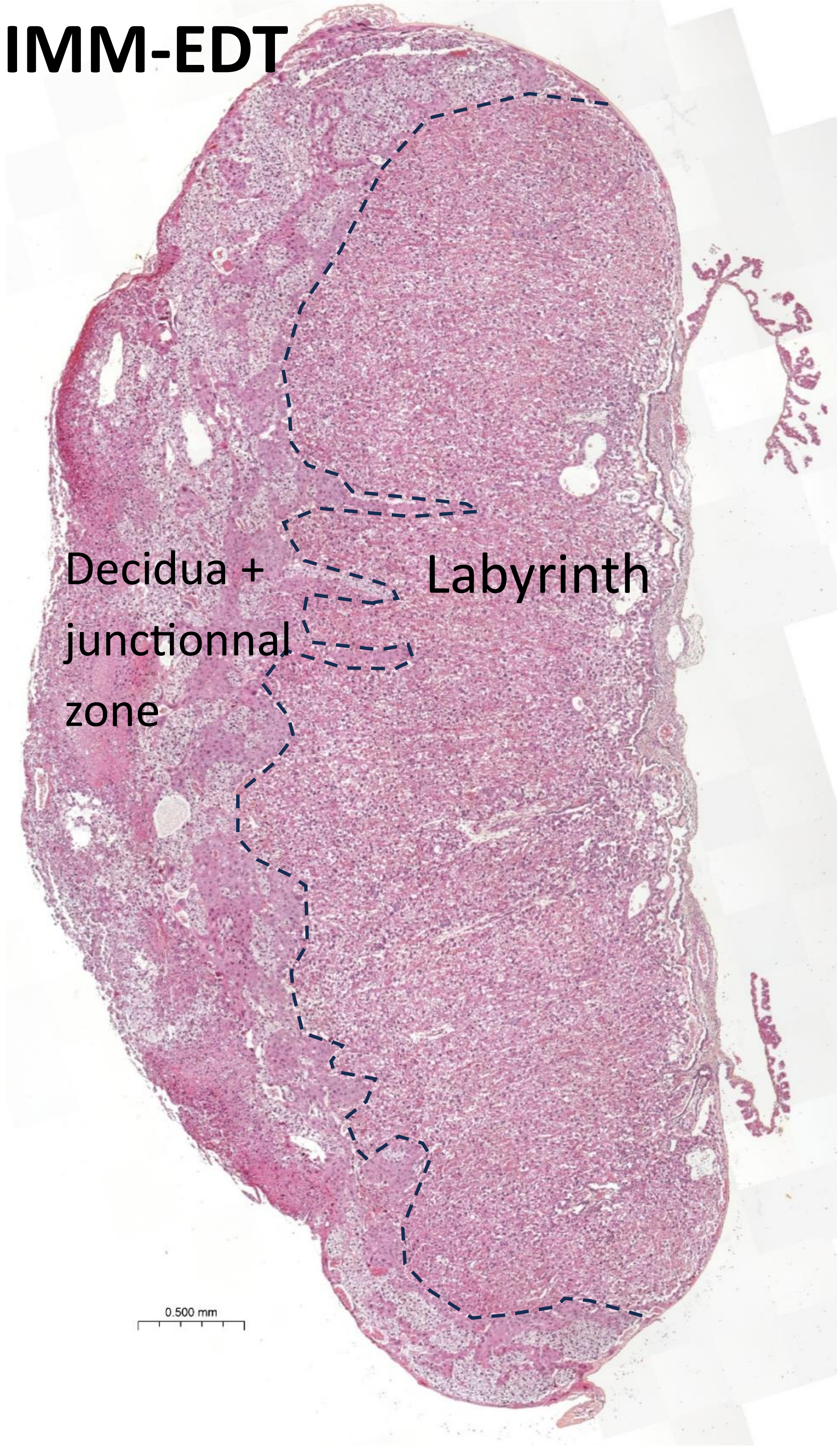


**Supplementary Figure 2: E18.5 placentas have the same global structures in Sham, PBS-EDT and IMM-EDT groups**

H&E staining of E18.5 placentas

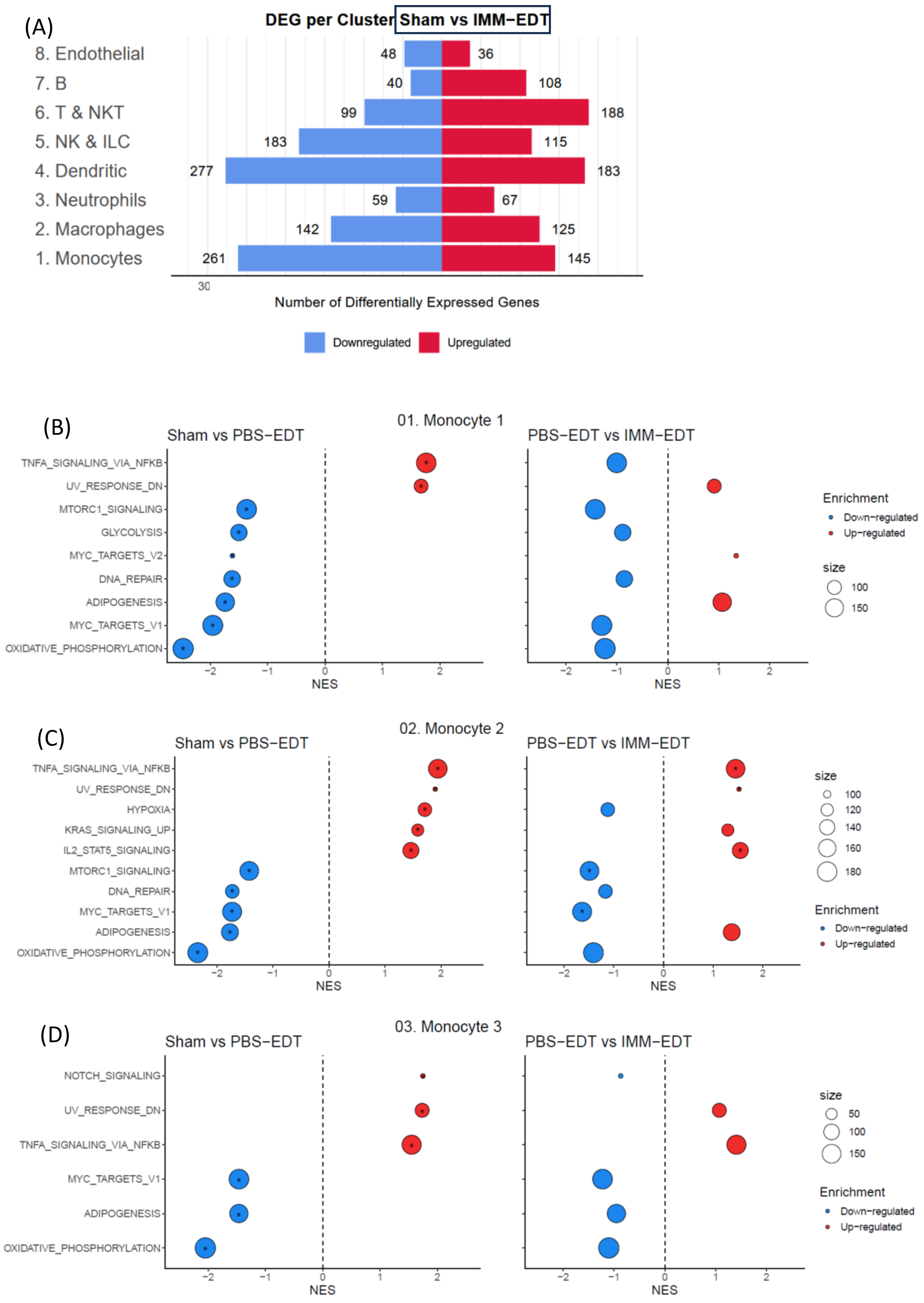
(A)
DEG per Cluster Sham vs IMM−EDT
8. Endothelial
7. B
6. T & NKT
5. NK & ILC
4. Dendritic
3. Neutrophils
2. Macrophages
1. Monocytes
48
36
40
108
99
188
183
115
277
183
59
67
142
125
261
145
30
Number of Differentially Expressed Genes
Downregulated
Upregulated
(B)
01. Monocyte 1
Sham vs PBS−EDT
PBS−EDT vs IMM−EDT
TNFA_SIGNALING_VIA_NFKB
UV_RESPONSE_DN
MTORC1_SIGNALING
GLYCOLYSIS
MYC_TARGETS_V2
DNA_REPAIR
ADIPOGENESIS
MYC_TARGETS_V1
OXIDATIVE_PHOSPHORYLATION
NES
Enrichment
Down−regulated
Up−regulated
size
100
150
(C)
02. Monocyte 2
Sham vs PBS−EDT
PBS−EDT vs IMM−EDT
TNFA_SIGNALING_VIA_NFKB
UV_RESPONSE_DN
HYPOXIA
KRAS_SIGNALING_UP
IL2_STAT5_SIGNALING
MTORC1_SIGNALING
DNA_REPAIR
MYC_TARGETS_V1
ADIPOGENESIS
OXIDATIVE_PHOSPHORYLATION
NES
size
100
120
140
160
180
Enrichment
Down−regulated
Up−regulated
(D)
03. Monocyte 3
Sham vs PBS−EDT
PBS−EDT vs IMM−EDT
NOTCH_SIGNALING
UV_RESPONSE_DN
TNFA_SIGNALING_VIA_NFKB
MYC_TARGETS_V1
ADIPOGENESIS
OXIDATIVE_PHOSPHORYLATION
NES
size
50
100
150
Enrichment
Down−regulated
Up−regulated

(E)

(F)

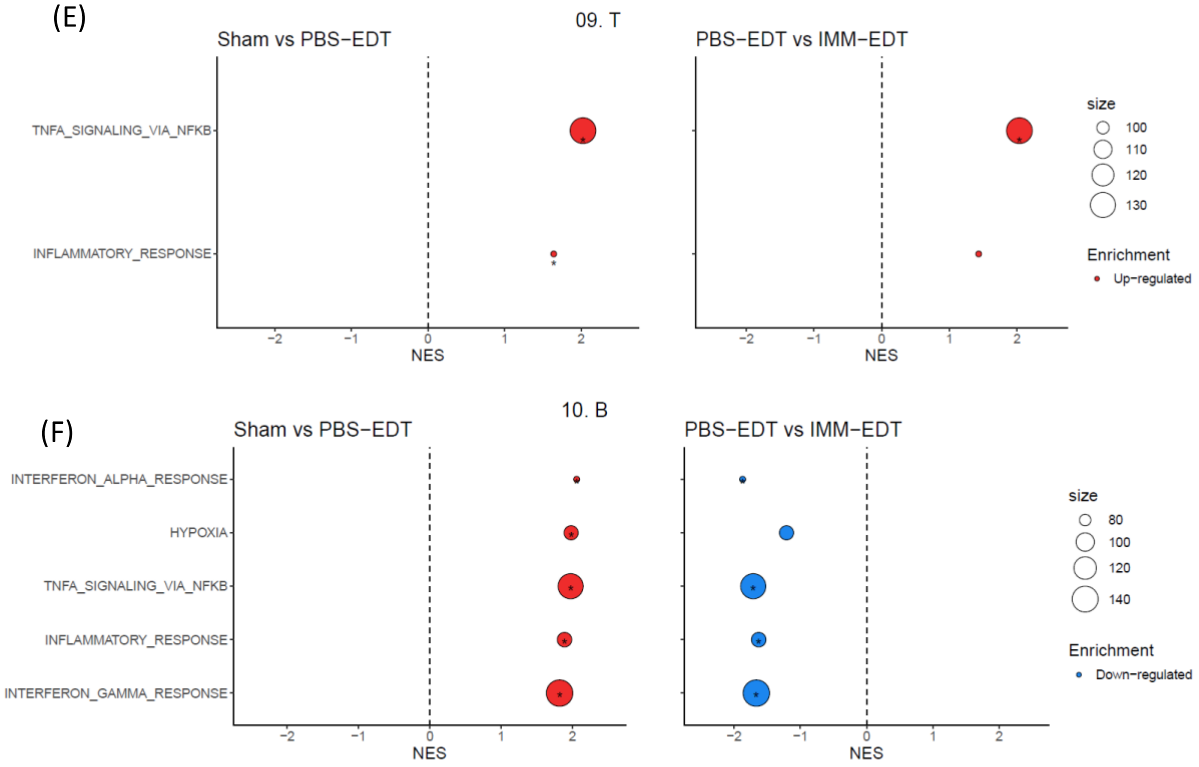


**Supplementary Figure 3: Differential expression analysis in immune cells reveals alterations driven by endometriosis and partially corrected by the IMM treatment** **(A)** Number of differentially expressed genes in each cell group for the groups Sham vs IMM-EDT. **(B)** Dotplot representing the top 10 enriched and negatively enriched pathways in Monocyte 1, **(C)** Monocyte 2, **(D)** Monocyte 3, **(E)** T cell and **(F)** B cells between Sham and PBS-EDT by Gene Set Enrichment Analysis.
